# Hypatia: Comparative Isoform Profiling Across Cell Populations from Long-Read Single-Cell Transcriptomes

**DOI:** 10.64898/2026.01.13.699341

**Authors:** Timothy Pan, Cheng-Kai Shiau, Lina Lu, Changhu Wang, Minhua Wang, Yueying He, Arvind Bhimaraj, Daniel Brat, Jason Huse, Jingyi Jessica Li, Ruli Gao

**Affiliations:** Department of Biochemistry and Molecular Genetics, Feinberg School of Medicine, Northwestern University, Chicago, IL, USA 60611; Center for Cancer Genomics, Robert H. Lurie Cancer Center, Northwestern University, Chicago, IL, USA 60611; Driskill Graduate Program, Northwestern University, Chicago, IL, USA 60611; Biostatistics Program, Public Health Sciences Division, Fred Hutch Cancer Center, Seattle, WA, USA 98109; Department of Biostatistics, University of Washington, Seattle, WA, USA 98109; Department of Cardiology, Methodist DeBakey Cardiology Associates, Houston Methodist Hospital, Houston, TX, USA-77030; Department of Pathology, Northwestern University – Feinberg School of Medicine, Chicago, IL, USA 60611; Department of Pathology and Translational Molecular Pathology, University of Texas MD Anderson Cancer Center, Houston, TX, USA 77030

**Keywords:** Isoform usage shift, Tsallis isoform diversity, Chi-square test

## Abstract

Long-read single-cell RNA-sequencing enables isoform-level quantification in individual cells, yet methods for delineating cell population-specific isoforms remain limited. Here, we develop Hypatia to systematically compare isoform complexities across three complementary dimensions, including isoform usage, diversity, and expression. Our analyses demonstrated that Cramer’s V improved the precision of detecting isoform usage shifts, at the cost of detecting fewer events in sparser data. Results further revealed Tsallis entropy as a refined, tunable measure of isoform diversity, improving comparability across genes and cells. Critically, integrating these analyses in real datasets provided complementary views of isoform variation across cell populations, highlighting Hypatia’s important applications.

## Background

Alternative splicing is an essential post-transcriptional mechanism that expands the coding capacity of eukaryotic genomes(1, 2). Despite its critical roles in health and disease, cell-type- and cell-state-specific RNA isoform complexity remains poorly understood due to the limitations of bulk and short-read sequencing technologies. More recently, long-read single-cell and single-nucleus RNA-sequencing (LR-scRNAseq)(3–8) achieves high-throughput and single-cell resolution of isoform quantification by coupling single-cell barcoding with long-read sequencing platforms (i.e., ONT and PacBio). However, a critical need remains for an integrated statistical framework that jointly evaluates isoform expression, diversity, and usage changes across cell populations, despite existing analytical frameworks that address important aspects of alternative splicing. Tools such as rMATS(9), DRIMSeq(10), DEXSeq(11), and diffSplice(12) focus on differential splicing events or transcript exon usage in replicated bulk RNA-sequencing data, while IsoformSwitchAnalyzeR(13) and tappAS(14) emphasize the functional interpretation of isoform changes but do not jointly address these three dimensions. TSIS(15) characterizes temporal isoform switches in bulk time-series data rather than variation across cell populations, and ScIsoX(16) quantifies multidimensional isoform diversity within and across cells but does not statistically test isoform usage shifts across cells. satuRn(17) provides scalable statistical testing of differential transcript usage in both bulk and single-cell RNA-sequencing but remains limited in scope, without assessing differential isoform expression or diversity. To address these gaps, we present Hypatia, a computational workflow for the integrated assessment and comparison of isoform expression, diversity, and usage as three complementary dimensions of isoform profiles across cell populations.

## Results and Discussion

Hypatia harnesses single-cell isoform expression data generated from LR-scRNAseq and performs three complementary modes of comparative analyses, evaluating 1) isoform usage, to identify differential isoform usage (DIU) events from changes in isoform fractions, 2) isoform diversity, to identify differential isoform diversity (DIV) events according to isoform species heterogeneity, and 3) isoform expression, to detect differentially expressed isoform (DEI) events across cell populations (**Fig. 1a**). Each analysis offers unique insights into cell-specific isoforms that are essential for understanding cellular function and disease mechanisms, integrating statistical methods for relative isoform usage shifts, species diversity, and expression abundance in a single platform. In addition, Hypatia leverages object-oriented R programming in the *SingleCellExperiment* class(18), which structures count data, cell-level and feature-level metadata in a synchronized manner, supporting seamless multi-scale data analysis.

**Figure 1.**
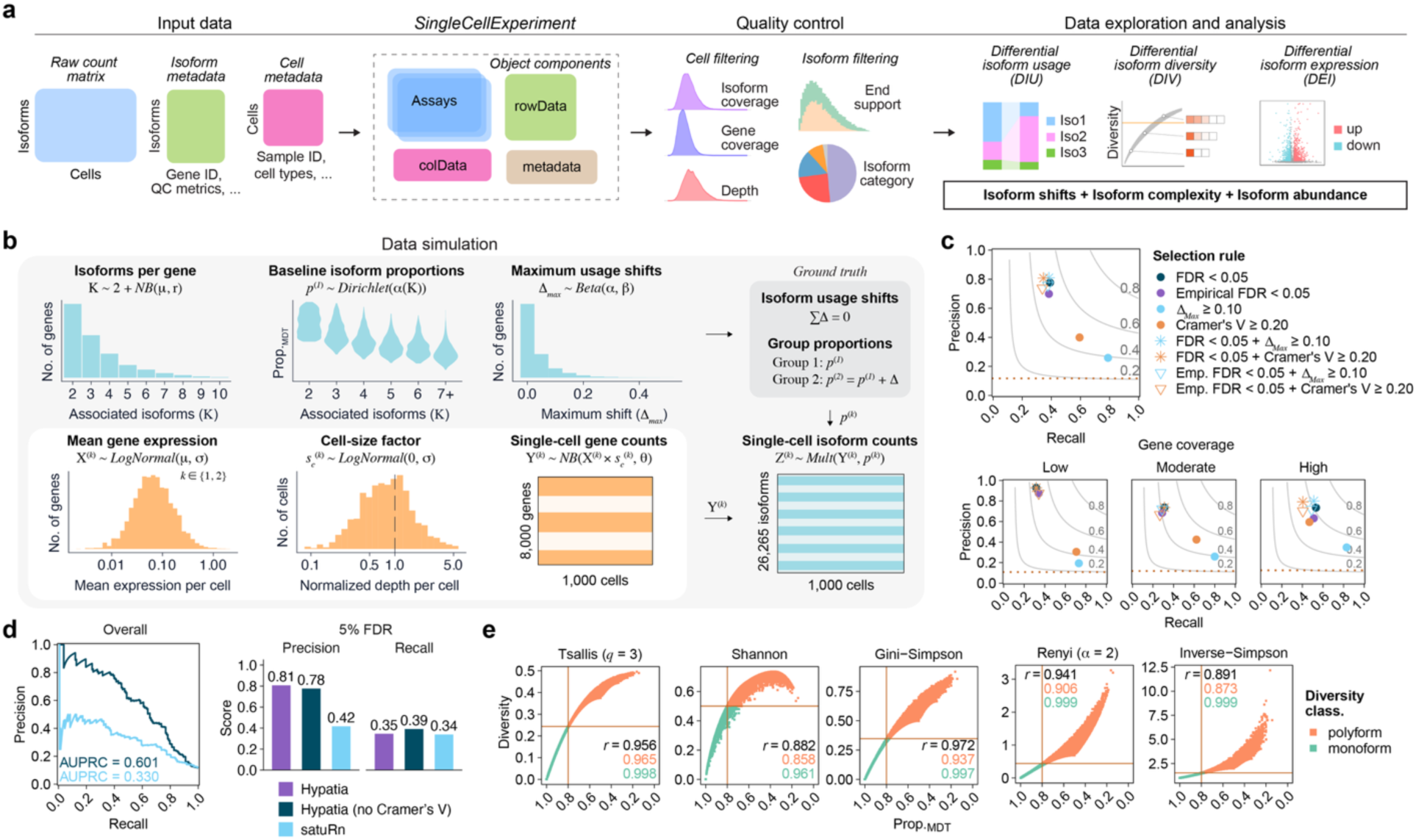
Hypatia workflow and simulation-guided assessment. a, Illustration of the main functional modules and workflow of Hypatia R package. **b,** Diagram outlining the steps in single-cell isoform count simulation. **c**, Precision-recall performance of DIU selection rules at overall and stratified gene coverage. Dashed horizontal line represents the ground-truth DIU prevalence. Contours represent F1 score. **d,** Comparison of DIU detection performance of Hypatia and satuRn. Left, precision-recall curves with the AUPRC values indicated. Hypatia’s DIU test was evaluated without filtering with Cramer’s V for fair comparison. Right, precision and recall at a nominal gene-level FDR < 0.05. **e,** Relationship between isoform diversity and proportion of the most dominant transcript (Prop_MDT_) of all simulated genes. Pearson correlation coefficients are shown for all data (black), polyform (orange), and monoform (turquoise) genes.

We evaluated statistical measurements via rigorous tests in simulation, where single-cell profiles were modeled using parameters derived from both in-house and published LR-scRNAseq datasets (**Fig. 1b, Table S1, Methods**). The simulated dataset closely recapitulated key characteristics of the real data, including cell-level sequencing depth and coverage, gene-level variation and abundance, and isoform-level complexity (**Fig. S1a-d**). With the simulated single-cell isoform data, we first evaluated DIU tests to establish robust criteria for detecting true isoform usage shifts between cell populations. The Chi-square test has been frequently applied in previous DIU analysis(19–21), yet its statistical reliability in the context of single cell data remains unclear. Our assessment revealed that Chi-square significance alone provided moderate performance in calling true DIU events, while permutation-based significance showed similar but slightly lower detection power (**Fig. 1c**). The filtering criterion, *Delta* ( Δ*_Max_* ), defined as the greatest difference in isoform proportions per gene between cell groups and the empirical analogue of the simulated effect used to define ground-truth DIU, tended to rescue recall with lower precision. Intriguingly, we found that the effect-size measure Cramer’s V (**Methods**) showed considerably better balance between precision and recall than the observed *Delta.* Furthermore, our analysis identified an optimal Cramer’s V cutoff of approximately 0.2 across all sequencing depth groups (**Fig. S2a-b**). While combining Chi-square significance with *Delta* provided acceptable power for DIU selection, incorporating Cramer’s V yielded modest but consistent improvements in precision, particularly for genes with higher coverage depth.

Next, we benchmarked Hypatia’s DIU test that couples Chi-square significance with Cramer’s V thresholding against a leading existing tool, satuRn, a quasi-binomial generalized linear modeling framework designed for both bulk and single-cell data(17). Notably, Hypatia demonstrated substantially better performance (AUPRC = 0.601) compared to satuRn (AUPRC = 0.330) (**Fig. 1d**, left). Consistently, Hypatia showed markedly higher precision than satuRn (0.81 versus 0.42), while maintaining comparable recall at a 5% FDR threshold (**Fig. 1d**, right). These data support Hypatia’s improved ability to detect true DIU events, although challenges inherent to single-cell data continue to limit achievable performance.

To complement isoform usage analysis, which detects significant differences in isoform usage proportions across cell populations, we further evaluated entropy-based methods for quantifying and comparing isoform diversity across genes and cellular groups. The widely used Shannon entropy(19, 20, 22) is sensitive to biases arising from gene-to-gene differences in isoform number and heavy-tailed abundance distributions associated with dropouts. Previous studies partially addressed this limitation by grouping isoforms into two abundance-based categories, thereby improving comparability across genes and reducing sensitivity to data noise (19, 20, 22). Here, we evaluated a range of entropy measures for their ability to quantify isoform diversity using the full distribution of all observed isoforms. Our analysis identified Tsallis entropy as a flexible measure of isoform diversity that adjusts the contribution of rare isoforms through a tunable parameter *q* (**Fig. 1e**). Analysis across a range of *q* values showed highest performance at *q* = 2 to 3, with *q* = 3 selected based on the slightly higher performance within the monoform and polyform strata (**Fig. S3**). Notably, comparison with other entropy measures revealed that Tsallis entropy (*q* = 3) demonstrated the tightest association with dominant transcript proportions, outperforming Shannon entropy and several other measures. Based on the approximately monotonic distribution of Tsallis isoform diversity scores, we applied thresholds to classify genes as monoform or polyform as well as their effective numbers of isoforms, providing categorical descriptions of isoform heterogeneity across genes. Moreover, we adopted bootstrap resampling to quantify within-group variation in Tsallis entropy and assess the statistical confidence of isoform diversity differences across cell groups. Together, these results support Tsallis entropy as a robust metric for quantifying and comparing isoform diversity across genes and cells, mitigating biases arising from differences in isoform number and dropout without requiring data subsetting.

To test Hypatia’s applications in profiling cell-type-specific isoforms in real-world data, we analyzed three LR-scRNAseq datasets, including an in-house primary glioblastoma (GBM02PR) dataset and two previously published datasets derived from a primary renal cell carcinoma (RCC12P)(3) and normal adult human heart tissue (HR182)(19) (**Table S1, Methods**). The DEI analysis revealed marked expression of unique isoforms associated with cell-type-specific genes in all three datasets, providing clear isoform-level resolution of cell-type-specific expression patterns (**Fig. 2a-c**). Next, applying Hypatia’s DIV analysis revealed pronounced differences in isoform heterogeneity across cell types (**Fig. 2d**). For instance, in RCC12P, *IGHA1* shifted from a polyform state expressing both membrane-bound and secreted isoforms in follicular B cells to a monoform state characterized by the dominant expression of the secreted isoform in plasma cells. Together, these analyses demonstrate that cell-type differences extend beyond isoform abundance to the overall complexity of isoform repertoires as captured by Hypatia’s DIV analysis.

**Figure 2.**
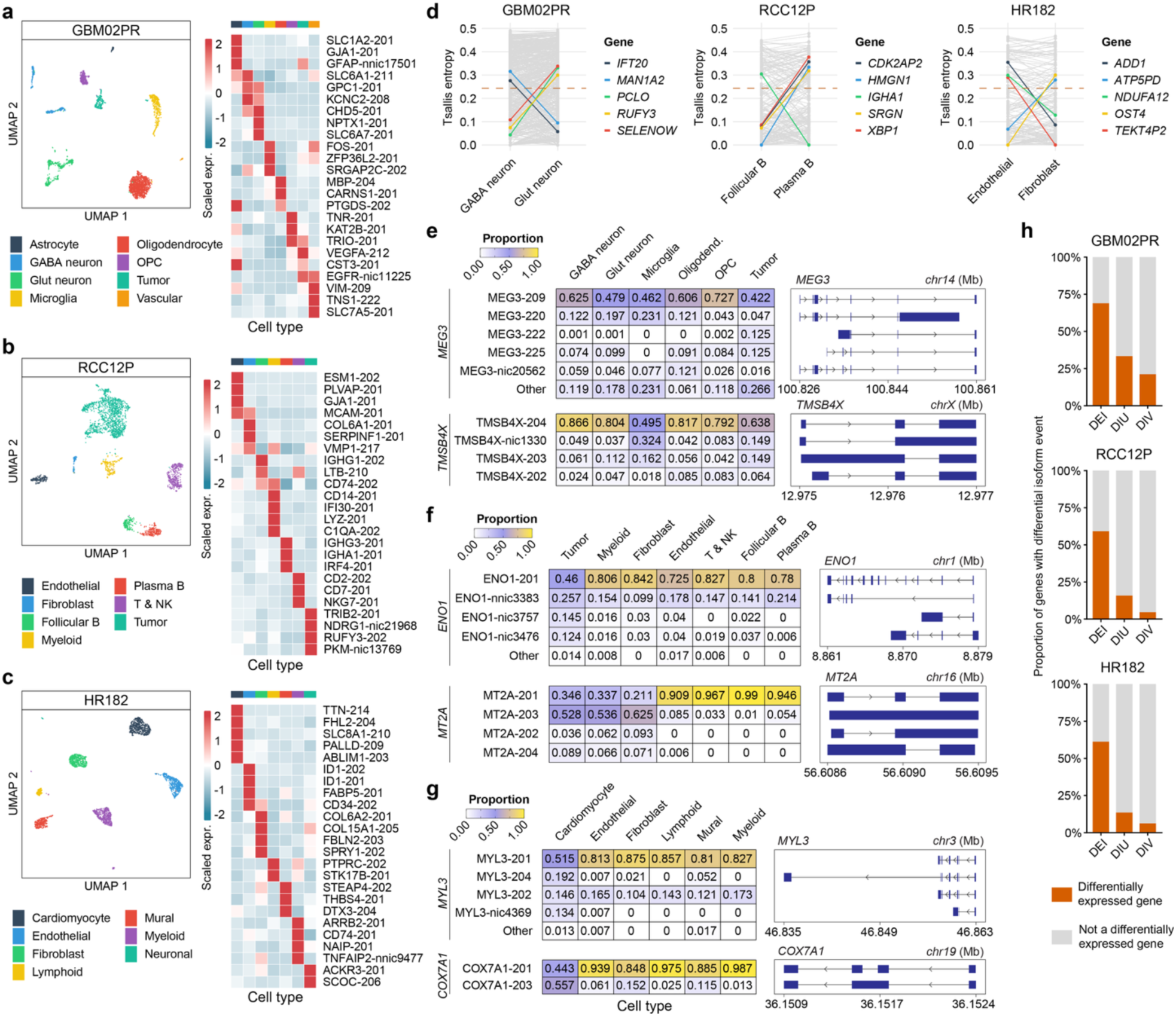
Hypatia characterizes isoform-level variations across cell-types in LR-scRNAseq. a-c, (left) UMAP determined by gene expression analysis and (right) heatmap of DEIs of each cell-type. **d,** Selected example genes exhibiting differential diversity between cell types in each sample. The horizontal dashed line indicates the entropy threshold used to classify monoform and polyform. **e-g,** (left) Heatmap showing cell-type-specific isoform proportions of DIU events in each sample. (right) Transcript structures of isoforms corresponding to heatmap rows. **h,** Stacked bar plots showing intersections between isoform-centric analyses and differential gene expression analyses.

Hypatia also identified extensive isoform usage shifts across cell-types, providing valuable insights into the regulatory mechanisms underlying cell-type-specific functions. In GBM02PR, multiple DIU events were identified in the tumor-suppressor lncRNA gene *MEG3* (**Fig. 2e**). The relative abundance of dominant isoform MEG3-209 (ENST00000451743) varied broadly across multiple cell-types, whereas MEG3-220 (ENST00000524035) and novel isoform MEG3-nic20562 had distinct expression in neoplastic tumor cells and oligodendrocytes, respectively. Similarly, the cytoskeleton-associated gene *TMSB4X* exhibited a marked microglia-specific decrease in protein-coding isoform TMSB4X-204 (ENST00000451311), accompanied by increased usage of the novel isoform TMSB4X-nic1330. Although the conserved actin binding motifs are preserved in this expanded novel isoform, it harbors a retained intron that is predicted to encode a disordered region within its open reading frame, warranting further investigation into the functional consequences of this isoform switch (**Fig. S4a**)(23). In RCC12P, significant DIU events were detected in glycolysis gene *ENO1* and metallothionein-encoding gene *MT2A* (**Fig. 2f**). Tumor cells expressed a distinct polyform state of *ENO1* characterized by uncatalogued isoforms, including ENO1-nnic3383, which is predicted to encode a truncated protein lacking the N-terminal domain important for dimerization and enzymatic function, potentially impairing α-enolase-mediated pathways in tumor cells (**Fig. S4b**)(23, 24). In a separate DIU event, a striking dichotomy in the dominant isoform of *MT2A* emerged across cell-types, recapitulating the previously reported isoform shift in RCC (3). Similarly, in the normal heart sample HR182, Hypatia recapitulated significant cardiomyocyte-specific DIU events in *MYL3* and *COX7A1*, key regulators of sarcomere contraction and energy production, respectively (**Fig. 2g**), consistent with previously reported isoform shifts(19). These findings demonstrate Hypatia’s utility in uncovering cell-type-specific splicing regulation of functionally relevant genes across diverse tissues.

We next examined the overlap among DIU, DIV, and DEI results to assess the complementary insights provided by these approaches. Among the 355 genes exhibiting changes in isoform complexities across cell types in GBM02PR, 93.5% were uniquely detected by only one of the three tests, including 224 genes by DEI, 68 genes by DIV and 40 genes by DIU. Similarly, up to 89.3% genes with cell type-specific isoform complexity in RCC and 80.4% in heart samples were uniquely revealed by only one test. Among genes detected by multiple analyses, DIU and DIV overlap accounted for 9 of 23 genes in GBM, 101 of 128 genes in RCC, and 54 of 77 genes in the heart sample (**Fig. S5**). These results exemplify the complementarity among the three types of isoform-centric analysis.

Lastly, we investigated their relationship to conventional gene-level differential expression analysis. Notably, DIV and DIU analyses revealed changes that were largely undetectable in conventional analysis for the differentially expressed genes (DEGs) across all three datasets, with only 13.6-33.4% DIU genes and 4.8-21.2% DIV genes overlapped with list of DEGs (**Fig. 2h**). By contrast, 59.2-68.9% DEI genes were also detected as DEGs, indicating potential confounding between the abundance-based analyses at isoform and gene levels. Nevertheless, DEI analysis provides additional resolution of cell-type-specific transcripts by identifying the specific isoforms contributing to overall gene expression profiles. These results demonstrate that isoform-level analyses offer a complementary and higher-resolution view of transcript regulation across diverse cell populations.

Collectively, our results highlight the utility of Hypatia for isoform-centric analysis of single-cell transcriptomes. We show that Tsallis entropy robustly quantifies isoform diversity and Cramer’s V effectively curates Chi-square statistics from long-read single-cell isoform data. Critically, we show that DEI, DIU and DIV tests are complementary in dissecting differential isoform patterns across cell populations. Results from patient-derived datasets show the prevalence of isoform-level variations across cell-types and exemplify the important applications of Hypatia in revealing comprehensive isoform spectrums of human tissues.

## Conclusions

In this study, we present Hypatia, an analytical framework for systematically dissecting isoform complexity from long-read derived single cell isoform data, extending transcriptomic profiling beyond gene-level expression to the comprehensive comparison of isoforms, the functional units of genes, across cell populations. By incorporating Tsallis entropy and Cramer’s V, Hypatia achieves reliable performance, although the technology-associated noise continues to constrain further improvements. Another limitation of Hypatia is its inability to discover previously unrecognized cellular subpopulations, while population-level aggregation may obscure multimodal or heterogeneous isoform patterns within a cell population. Lastly, integrating Hypatia with complementary tools, such as acorde(25) for isoform co-regulation network analysis and tappAS(14) and IsoformSwitchAnalyzeR(13) for functional interpretation, could connect population-specific isoform changes to their regulatory relationships and functional consequences, providing broader insights into isoform biology.

## Methods

### Simulation of single-cell isoform expression data

To evaluate method performance under controlled conditions, single-cell isoform expression data were simulated to reflect empirical characteristics of long-read transcriptomic data. Parameters were estimated from quality-filtered long-read single-nucleus RNA-sequencing datasets described below, restricted to genes with at least two observed isoforms. Isoform number was modeled using a shifted negative binomial distribution (offset = 2, size = 1.092, *μ* = 1.291), while gene abundance and cell-size factors were modeled independently using log-normal distributions. Baseline isoform proportions were modeled using symmetric Dirichlet distributions for genes with 2, 3, 4, 5, or ≥6 isoforms. The corresponding Dirichlet parameters (α = 1.122, 1.000, 1.000, 1.000, and 0.794) were selected by minimizing differences in dominant isoform proportions and Shannon entropy between observed and simulated data. Using these fitted distributions, we simulated 8,000 genes in two populations of 500 cells each. Baseline isoform proportions were drawn from the corresponding Dirichlet distribution, and maximum usage differences were drawn from a beta distribution (*α* = 0.5, *β* = 10). 25% of genes were designated exact nulls. The remaining isoform usage difference (Δ) vectors were generated and constrained for valid group-specific proportions. Group-specific gene abundances were drawn independently from a cell-number-adjusted log-normal distribution (meanlog = -2.956, sdlog = 1.126), and cell-size factors were drawn from a log-normal distribution (meanlog = 0, sdlog = 0.687) and normalized within each population. Per-cell gene counts were sampled from a negative binomial distribution (size = 0.10) and allocated among isoforms using the group-specific proportions and a multinomial distribution.

### Differential isoform usage performance and benchmarking

To evaluate DIU performance, genes with a maximum absolute isoform usage difference (Δ*_Max_*) ≥0.10 were designated as ground-truth DIU events. Precision and recall were calculated using genes with at least two retained isoforms, detected in ≥5% of cells per group, and supported by ≥15 total counts per group. The Chi-square and empirical P-values were calculated following the isoform usage analysis section below. For coverage-stratified analysis, genes were divided into tertiles based on total counts, corresponding to low (67-132), moderate (133-205), and high (206-2,138) coverage groups.

To benchmark Hypatia against satuRn(17) (v1.18.0), the simulated count matrix was analyzed using Hypatia’s RunDIU() function and satuRn’s fitDTU() and testDTU() functions with a ∼0 + group design. Performance was evaluated on the same filtered gene set. Precision-recall curves ranked genes by Hypatia’s Chi-square P-value or the minimum empirical transcript-level P-value reported by satuRn. At a nominal 5% FDR, Hypatia’s calls were based on Benjamini-Hochberg-adjusted P-values with or without Cramer’s V ≥0.20, whereas satuRn calls were based on the recommended DEXSeq(11) (v1.56.0) gene-screening q-values derived from empirical transcript-level P-values.

### Sample acquisition and LR-snRNAseq

Fresh-frozen tissue from a grade 4 glioblastoma of the frontal lobe was collected from the NBST tissue bank at Northwestern University and subjected to snNanoRNAseq(3). The single-nuclei suspension was prepared using the Chromium Nuclei Isolation Kit (10X Genomics) following the manufacturer’s protocol. Nuclei were then loaded onto a 10X Genomics Chromium Controller (iX) with Chip J with an expected recovery of 3,000 nuclei. Full-length barcoded cDNAs were amplified and enriched for long fragments. The final library was prepared using the SQK-LSK114 ligation sequencing kit (Oxford Nanopore Technologies) and sequenced on an in-house PromethION with an R10.4M flow cell.

### LR-snRNAseq data processing

Raw single-nucleus sequencing data of the GBM02PR, RCC12P and HR182 samples were processed using scNanoGPS(3) (version 2.0). Isoform quantification was performed with IsoQuant(26) (version 3.4.2) using the ‘unique only’ transcript quantification setting and reference genome GRCh38 (GENCODE v44/Ensembl110 annotations). Reads mapping to *MALAT1* and *NEAT1* were excluded due to computational restraints. SQANTI3(27) (version 5.2.2) was used for transcript annotation. Transcripts categorized as full-splice match were retained. Transcripts not present in the reference annotation, classified as novel-in-catalog (nic) and novel-not-in-catalog (nnic), required additional support of 5’ CAGE peaks in refTSS(28) (version 4.1) and 3’ polyA sites in PolyASite(29) (version 3.0). All transcripts were required to be detected in at least 3 cells in the sample and free of predicted RT-switching artifacts.

### Single-cell gene expression analysis

Single-cell gene expression analysis was performed using Seurat(30) (version 5.0.1). Cells with expression of more than 200 genes and less than 5% of mitochondrial genes were retained. Clustering utilized the top 15 principal components and cell-types were annotated using established gene markers(31). Significant differentially expressed genes (DEGs) were those that had |Log2 (fold change)| ≥0.60, adjusted *P*-value ≤0.05, and detected in ≥5% of cells in comparisons.

### Isoform usage analysis

DIU analysis of real datasets compared each cell-type against all other cell-types. DIU events were determined by Hypatia’s ‘RunDIU’ function with default settings, testing genes expressed in ≥5% of cells and supported by ≥15 counts in each group compared. A contingency table was constructed for each gene using aggregated isoform counts of each cell group being compared. Chi-square P-values were calculated using Pearson’s Chi-square test without continuity correction and adjusted across the evaluated genes using the Benjamini–Hochberg procedure to control the FDR. Bootstrapped 95% confidence intervals for isoform proportion differences were estimated using 5,000 iterations with cells resampled with replacement to original sample sizes. For performance evaluation, empirical p-values were calculated using the Pearson Chi-square statistic from 10,000 cell-label permutations that preserved group sizes, then adjusted across the evaluated genes to obtain Benjamini-Hochberg FDRs.

Effect size was calculated as Cramer’s V (**Eq. 1**)(32),

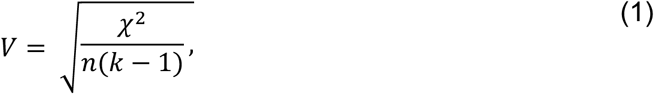

where *χ*^2^ denotes the Chi-square statistic, *n* denotes the total isoform count of the gene, and *k* is the number of columns in the contingency table (*k* = 2 for two-group comparisons). DIU events with an Benjamini-Hochberg-adjusted P-value ≤0.05 and Cramer’s V ≥0.20 were considered significant.

### Isoform diversity analysis

Isoform diversity was calculated using Tsallis (**Eq. 2**, *q* = 3), Shannon (**Eq. 3**), Gini-Simpson (**Eq. 4** or **Eq. 2** with *q* = 2), Renyi (**Eq. 5**, α = 2), and Inverse-Simpson (**Eq. 6**) entropies as implemented in the ‘RunDIV’ function of Hypatia. The entropies were calculated as the following,

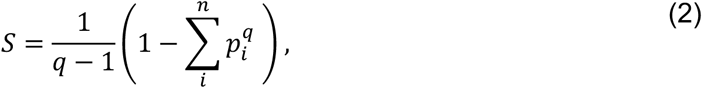

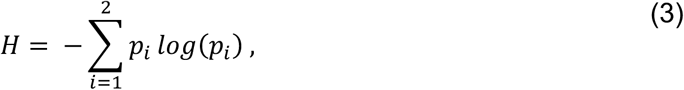

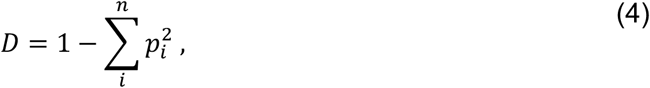

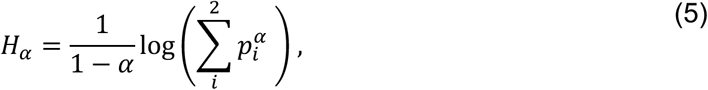

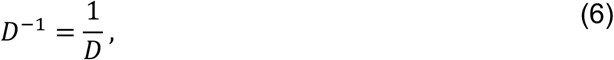

where *n* denotes the total number of expressed isoforms of the gene, *p* is the isoform proportion, and *q* and *α* represent the order of the Tsallis and Renyi entropies, respectively. The Shannon and Renyi entropies were restricted to the top 2 isoform abundances to improve comparability across genes and mitigate distinct strata driven by isoform number. Cutoffs for monoform and polyform classifications were guided by a Prop_MDT_ = 0.80, which corresponded to approximate limiting entropies at 0.243 (Tsallis), 0.500 (Shannon), 0.348 (Gini-Simpson), 0.435 (Renyi), and 1.533 (inverse-Simpson).

DIV events were assessed using Hypatia’s ‘RunDIV’ function with default settings, where genes required ≥5% detection and ≥15 supporting isoform counts in each group being compared. For bootstrapping, cells were sampled with replacement in each group across 5,000 iterations, and isoform diversity was calculated with **Eq. 2** (*q* = 3). Cells were sampled with replacement within each group to match original sample sizes across 5,000 iterations. Isoform counts were aggregated, and diversity was calculated using Eq. 2 (q = 3). Bootstrap diversity differences were summarized by their mean and 95% percentile intervals. Significant DIV events were considered bootstrap-supported when ≥97.5% of valid bootstrap differences were ≥0.10 or ≥97.5% of valid bootstrap differences were ≤-0.10.

### Isoform expression analysis

Isoform count data were first normalized using ‘NormalizeCounts’ in Hypatia with the ‘LogNormalize’ method, which divides isoform counts by the total counts in each cell, multiplies the resulting values by a scale factor of 10,000, and applies a natural log transformation. DEI events were calculated using ‘RunDEI’, which applies the Wilcoxon Rank Sum Test with Bonferroni corrected *P*-values. Significant DEIs were those that had |Log2 (fold change)| ≥0.60, adjusted *P*-value ≤0.05, and detected in ≥5% of cells in comparisons.

### Overlap of differential isoform events and differentially expressed genes

Genes presenting significant differential isoform events and gene expression were enumerated per cell-type comparison, where each cell-type was compared against all others. Overlap proportions were calculated within each cell type and subsequently averaged across cell types.

## Declarations

### Ethics approval and consent to participate

The IRB protocol for studying the multi-omics profiles of human tissues was approved by Northwestern University (STU00219936). Consent was waived because this study only involves the secondary analysis of the banked tissues, which has no direct interaction with subjects.

## Consent for publication

Not applicable.

## Availability of data and materials

Both raw and processed data generated in this study have been deposited in the Gene Expression Omnibus (GEO) database under accession GSE310974. Previously published sequencing data were downloaded from SRA under runs <u>SRR21492157</u> (RCC) and <u>SRR32154432</u> (heart). Simulation data have been deposited in Zenodo (doi: 10.5281/zenodo.17781226). All source code and documentation of Hypatia are open source and are released to the public and freely available on GitHub at https://github.com/gaolabtools/Hypatia.

## Authors’ contributions

TP led the development of Hypatia, performed data analysis, and wrote the manuscript. CKS processed raw sequencing data and reviewed results. MW performed the sequencing experiment and edited the manuscript. LL and YH reviewed results. CW and JJL participated in data analysis, reviewed results, and edited the manuscript. JH, DB and AB assisted with sample collection and edited the manuscript. RG supervised the study, conceived concepts, designed the computational tool, and wrote the manuscript.

## Acknowledgements

We thank all our funding supports and patients who consented for research use of their tissue samples.

