## Supplemental Figures for "Hypatia: Comparative Isoform Profiling Across Cell Populations from Long-Read Single-Cell Transcriptomes"

**Pan *et al*., 2026**

### Supplemental Material

### Figures S1-S5

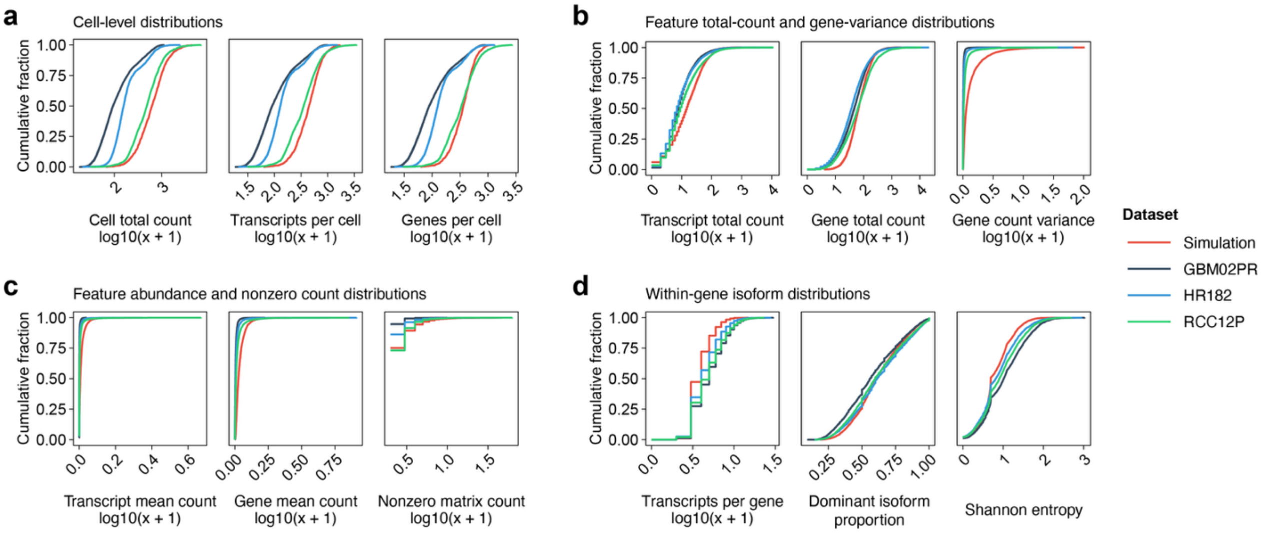


**Figure S1. Characteristics of simulated and real long-read single-cell RNA sequencing datasets.** Cumulative distributions of **a,** Cell-level features, including the total counts, detected transcripts and genes per cell; **b,** total transcript and gene counts, and gene-level count variance across cells; **c,** average transcript and gene counts, and nonzero entries in isoform matrix; and **d,** the number of observed isoforms per gene, proportion of the most dominant isoform, and gene-level Shannon entropy of isoform usage. Count matrices were restricted to genes with at least two isoforms and curves represent up to 5,000 randomly sampled finite observations per dataset and metric. The real datasets included glioblastoma (GBM02PR), renal cell carcinoma (RCC12P) and heart without disease conditions (HR182).

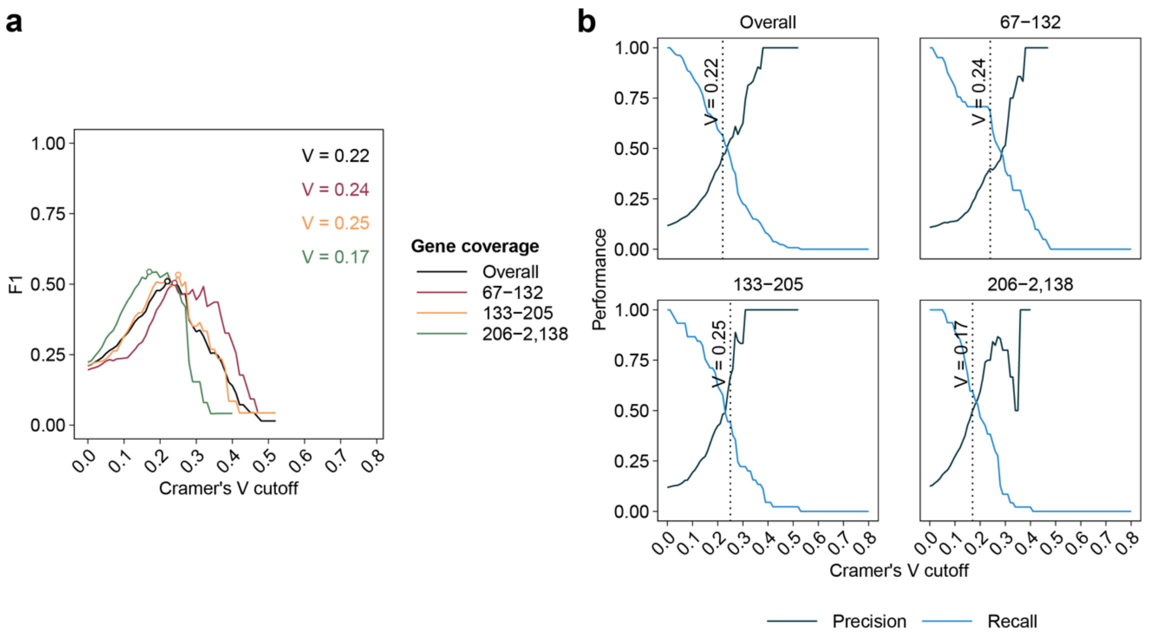


**Figure S2. Selection of the Cramer's V effect size cutoff for differential isoform usage analysis. a,** F1 score across Cramer's V thresholds for all simulated genes and each gene-coverage group. Circles indicate the cutoff that maximized F1. Lowest values were selected in the event of a tie. The final cutoff was set at 0.20 based on the approximate average of the estimated optimal thresholds. **b,** Corresponding precision-recall trade-offs for each coverage stratum. Dotted vertical lines indicate the F1-optimal cutoff for each gene-coverage group.

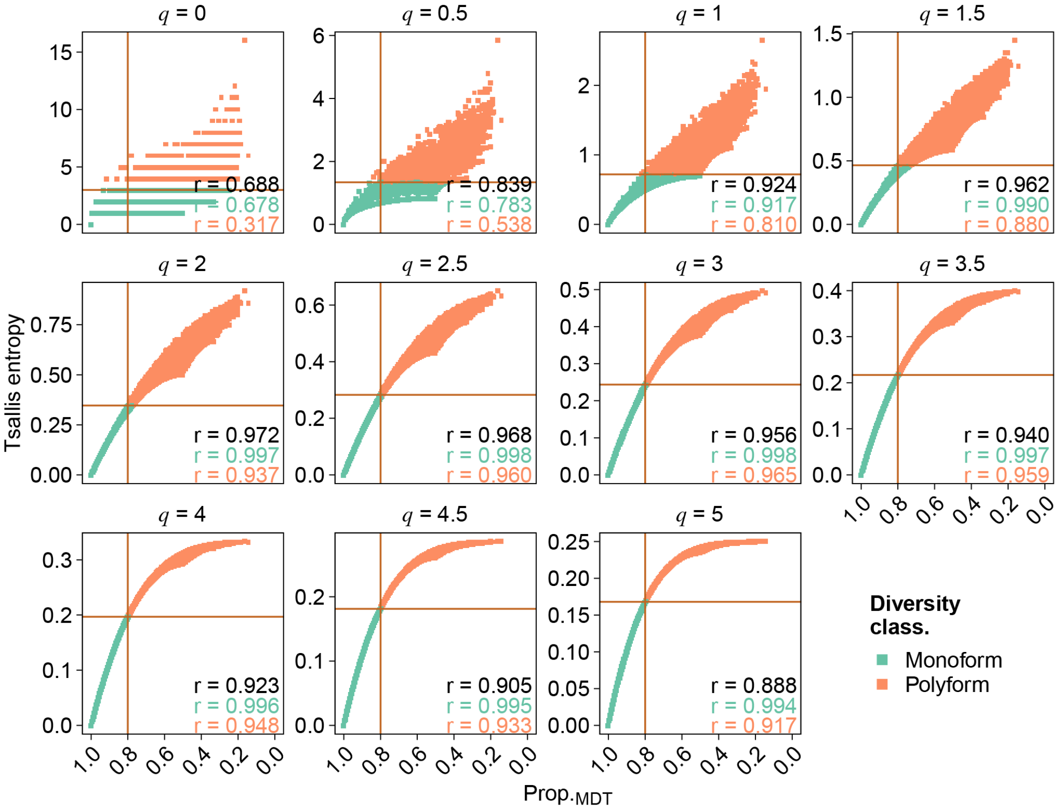


**Figure S3. Sensitivity of isoform-diversity classification to Tsallis entropy order. a,** Tsallis entropy plotted against the proportion of the most dominant transcript ($Prop._{MDT}$) on all simulated genes across orders ($q$ = 0 to $q$ = 5 in increments of 0.5). The vertical line marks Prop.MDT = 0.80 and the horizontal line denotes the corresponding order-specific entropy cutoff. Profiles at or below the cutoff were classified as monoform, whereas above were classified as polyform. Pearson's correlation of overall and each diversity class are annotated.

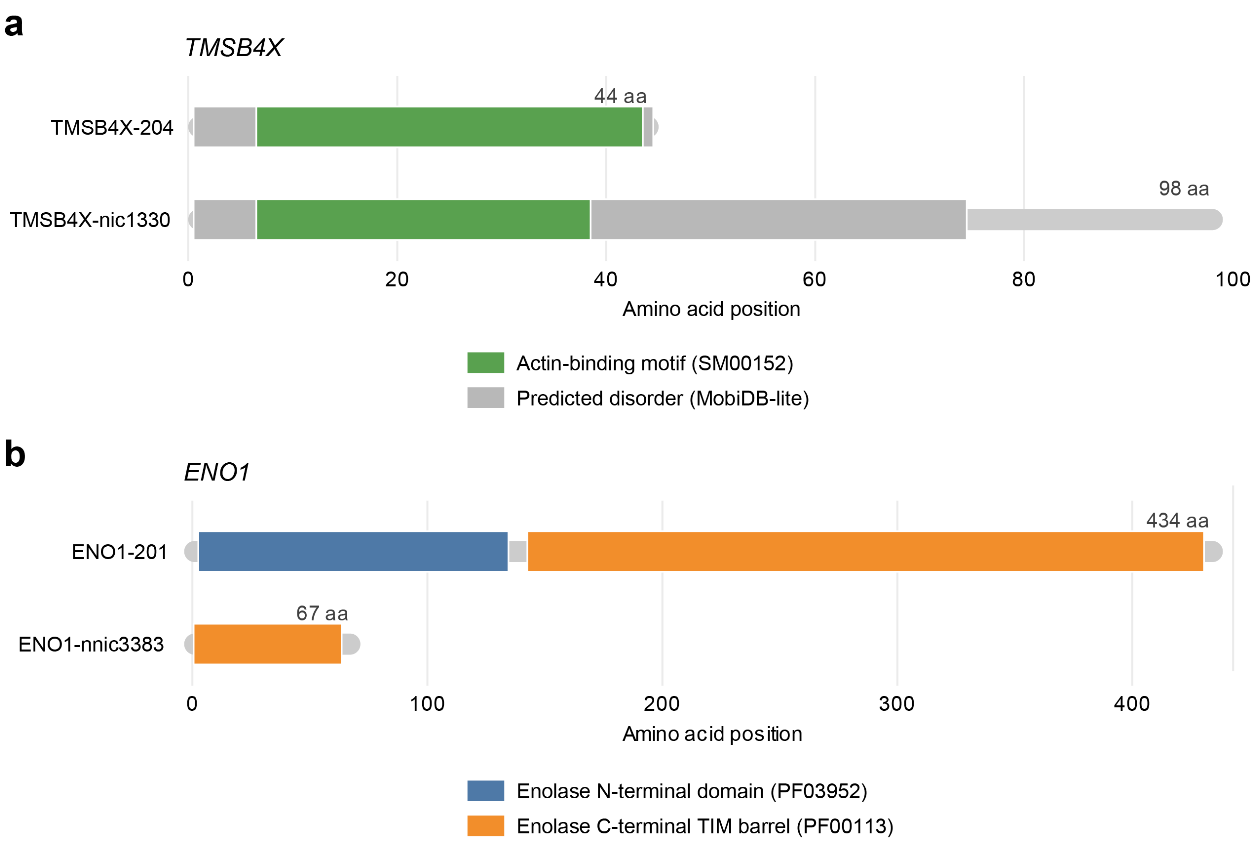


**Figure S4. Predicted protein domain organization of *TMSB4X* and *ENO1* isoforms.** InterProScan 6 annotations of canonical and novel (**a**) *TMSB4X* and (**b**) *ENO1* isoforms plotted to amino-acid scale. Representative Pfam and SMART annotations and the longest MobiDB-lite interval per isoform are shown.

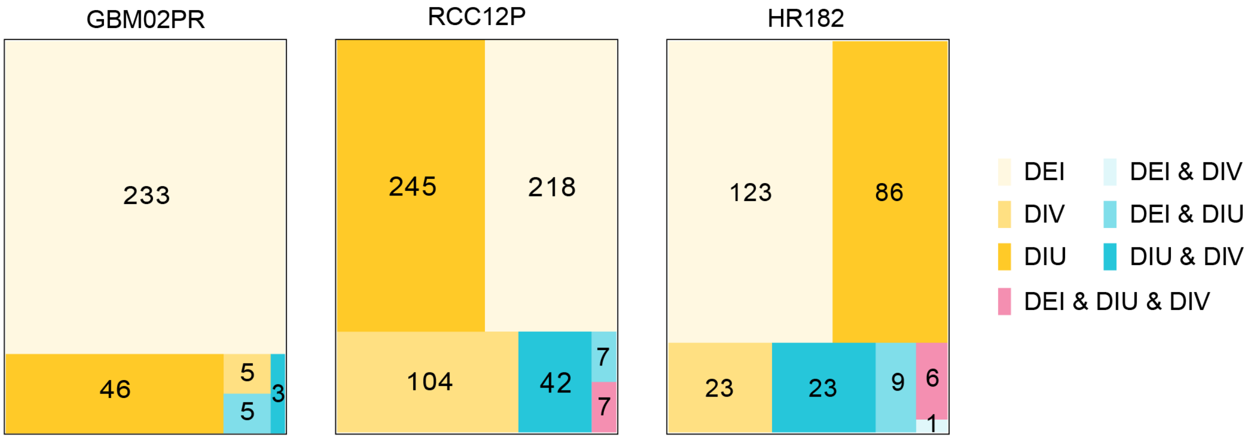


**Figure S5. Overlap of DIU, DIV, and DEI events.** Treemaps of DIU, DIV, and DEI genes across cell types in primary glioblastoma (GBM02PR), primary renal cell carcinoma (RCC12P) and non-diseased heart (HR182) datasets, shaded by unique and overlapping event type categories.
